# The Scm FCS finger binds the Pcl Tudor domain to link canonical PRC1 and PRC2.1 in Drosophila

**DOI:** 10.64898/2026.09.23.753712

**Authors:** Berfin Yesil, Barbara Steigenberger, Jacques Bonnet, Jürg Müller

## Abstract

Canonical Polycomb Repressive Complex 1 (PRC1) was previously reported to selectively associate with Polycomb Repressive Complex 2.1 (PRC2.1) in cross-linking-based affinity purification experiments from *Drosophila* embryos, but the molecular basis of this interaction has remained unresolved. Here, we established an affinity-purification strategy using cryo-milled *Drosophila* embryos that recovers intact canonical PRC1 together with the complete PRC2.1 complex without chemical cross-linking. Using AlphaFold3 to systematically model interactions between canonical PRC1 and PRC2.1 subunits, we identified a single high-confidence interface between the FCS finger of the canonical PRC1 accessory subunit Sex comb on midleg (Scm) and the atypical Tudor domain of the PRC2.1 accessory subunit Polycomblike (Pcl). Recombinant Scm FCS finger and Pcl Tudor domain formed a stable complex in vitro, as demonstrated by size-exclusion chromatography and native mass spectrometry. Structure-guided mutagenesis validated the predicted interface: point mutations in either interaction partner abolished complex formation, whereas complementary charge-reversal mutations restored binding. These findings identify the Scm–Pcl interaction as the molecular basis for the selective association between canonical PRC1 and PRC2.1 in *Drosophila* and reveal that communication between these complexes is mediated by a direct interaction between these accessory subunits.

## INTRODUCTION

In *Drosophila*, Polycomb group (PcG) protein complexes assemble at Polycomb Response Elements (PREs) to establish and maintain transcriptional repression of associated target genes. PREs are co-occupied by the sequence-specific DNA-binding Pho Repressive Complex (PhoRC), canonical Polycomb Repressive Complex 1 (PRC1), Polycomb Repressive Complex 1.1 (PRC2.1), and the Polycomb Repressive Deubiquitinase complex (PR-DUB) at hundreds of target genes in embryos and larvae (Kang et al., 2022; Klymenko et al., 2006; Nekrasov et al., 2007; Oktaba et al., 2008; Papp & Müller, 2006; Scheuermann et al., 2010; Schuettengruber et al., 2009; Schwartz et al., 2006). Although several sequence-specific DNA-binding proteins have been proposed to contribute to Polycomb complex recruitment to PREs (Brown et al., 1998; Brown et al., 2018; Klymenko et al., 2006; Ray et al., 2016), the molecular mechanisms are best understood for Pho, the sequence-specific DNA-binding subunit of PhoRC and the *Drosophila* ortholog of mammalian YY1. In PhoRC, Pho recruits the chromatin protein Sfmbt to PRE DNA. Genetic, biochemical, and structural studies established that Sfmbt binds the canonical PRC1 accessory subunit Sex comb on midleg (Scm), which in turn interacts with the PRC1 core subunit Polyhomeotic (Ph) through ordered SAM-domain interactions, providing the molecular basis for anchoring canonical PRC1 to PRE-bound PhoRC (Frey et al., 2016; Kim et al., 2005). Consistent with this model, mutation of Pho-binding sites in PRE reporter genes abolishes PhoRC binding and canonical PRC1 occupancy (Brown et al., 2023; Frey et al., 2016). Importantly, mutation of the same Pho-binding sites also markedly reduces PRC2 occupancy and H3K27me3 domain formation (Brown et al., 2023; Frey et al., 2016). Likewise, H3K27me3 deposition at Polycomb target genes is compromised in *pho* mutant animals (Brown et al., 2018). Together, these findings indicate that PhoRC coordinates the recruitment or stable association of both canonical PRC1 and PRC2.1 at PREs.

Although these studies established the molecular basis for recruitment of canonical PRC1 by PhoRC, they did not explain how PRC2.1 is physically connected to PRE-bound PhoRC or canonical PRC1. Cross-linking-based affinity purification experiments subsequently identified a selective association between canonical PRC1 and PRC2.1 in *Drosophila* embryos and implicated Scm as the factor linking the two complexes (Kang et al., 2022; Kang et al., 2015). Independent genetic evidence also supported a functional connection between Scm and PRC2: during differentiation of nurse cells in the *Drosophila* female germline, Scm was required for efficient PRC2-dependent H3K27me3 domain formation at Polycomb target genes (DeLuca et al., 2020). Together, these findings implicated Scm in the physical association between canonical PRC1 and PRC2.1 but left the molecular basis of this interaction unresolved.

Here, we established an affinity-purification strategy that enabled isolation of intact canonical PRC1 together with the complete PRC2.1 complex from *Drosophila* embryos under native conditions. Using this approach, we identify a direct interaction between the Scm FCS finger and the atypical Tudor domain of the PRC2.1 accessory subunit Polycomblike (Pcl) and show that this interaction provides the molecular basis for the selective association between canonical PRC1 and PRC2.1.

## RESULTS

### Affinity purification of Polycomb complexes from cryo-milled embryos preserves higher-order Polycomb assemblies under native conditions

Previous biochemical purifications of *Drosophila* Polycomb protein complexes relied on affinity purification of genetically tagged Polycomb proteins from soluble nuclear extracts. In these preparations, nuclei were extracted with high salt to solubilize nucleoplasmic and chromatin-associated proteins, and insoluble chromatin was subsequently removed. This approach enabled the isolation of canonical PRC1, PRC2.1, PhoRC, and PR-DUB as stable biochemical entities with well-defined subunit compositions (Czermin et al., 2002; Klymenko et al., 2006; Müller et al., 2002; Nekrasov et al., 2007; Scheuermann et al., 2010; Shao et al., 1999). However, because high-salt extraction dissociates many protein–protein interactions and separates protein complexes from chromatin, it is expected to disrupt higher-order assemblies that may exist when multiple Polycomb complexes co-occupy PREs. Consequently, interactions between Polycomb complexes were generally not detected or were recovered only as substoichiometric associations (Frey et al., 2016).

To recover such assemblies under native conditions, we developed an affinity-purification strategy based on cryo-milling of *Drosophila* embryos followed by low-salt extraction and affinity purification (**Figure 1A**). Because chromatin is mechanically disrupted before extraction, this procedure was designed to recover not only soluble Polycomb complexes but also complexes associated with fragmented chromatin, thereby increasing the likelihood of preserving higher-order Polycomb assemblies.

**FIGURE 1.**
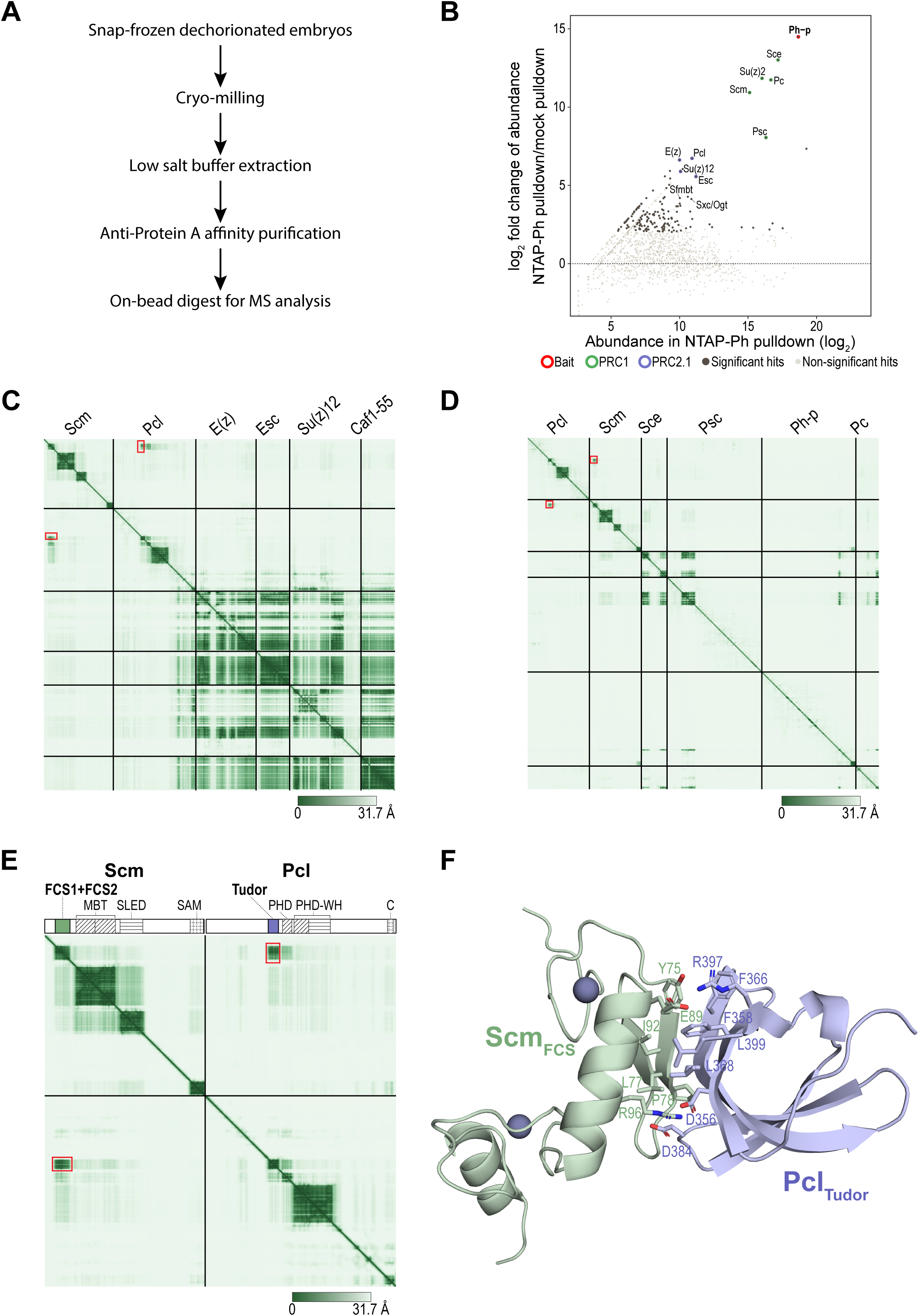
Native isolation of canonical PRC1–PRC2.1 assemblies and identification of the Scm–Pcl interaction by AlphaFold3. **A**: Workflow for affinity purification of NTAP-Ph from cryo-milled *Drosophila* embryos. **B**: Mass spectrometric analysis of proteins enriched in the NTAP-Ph affinity purification compared with the mock control purification from wild-type Oregon-R embryos. Identified PRC1, PRC2.1, and PhoRC subunit Sfmbt and Sxc/Ogt are indicated. **C-E**: AlphaFold3 analysis of interactions between canonical PRC1 and PRC2.1 subunits, identifying the Scm FCS finger–Pcl Tudor domain interaction as the only high-confidence predicted interaction. **C:** Predicted Aligned Error (PAE) plot (Elfmann & Stülke, 2023) for Scm and the five PRC2.1 subunits. **D:** PAE plot for Pcl, Scm, and core subunits of canonical PRC1. **E:** PAE plot for Scm and Pcl. **F**: AlphaFold3 model of the Scm FCS finger–Pcl Tudor domain complex. Residues forming the predicted hydrophobic and electrostatic interactions are shown as sticks. See Figure S1 for a superposition of the five AlphaFold3 models.

Using a previously described transgenic line expressing Protein A-tagged Ph, we affinity-purified associated proteins from cryo-milled *Drosophila* embryos under native conditions. In addition to the complete canonical PRC1 complex together with its accessory subunit Scm, the purification recovered the PhoRC subunit Sfmbt, the PRC2.1 subunits Pcl, E(z), Su(z)12, and Esc, and the O-GlcNAc transferase Sxc/Ogt (**Figure 1B**) (Frey et al., 2016; Gambetta & Müller, 2014; Gambetta et al., 2009; Nekrasov et al., 2007).

The co-isolation of PRC2.1 with canonical PRC1 under native conditions, together with the proposed role of Scm in linking the two complexes identified in cross-linking-based affinity purification experiments (Kang et al., 2015), prompted us to investigate the molecular basis of this interaction.

### AlphaFold3 identifies the Scm FCS finger–Pcl Tudor interface as the only high-confidence predicted interaction between canonical PRC1 and PRC2.1

We used AlphaFold3 (Abramson et al., 2024) to systematically model interactions between canonical PRC1 and PRC2.1 subunits. Among all combinations, AlphaFold3 identified a single high-confidence interaction between the FCS finger of the PRC1 accessory subunit Scm and the Tudor domain of the PRC2.1 accessory subunit Pcl, whereas no interactions were predicted between Scm and PRC2 core subunits or between Pcl and PRC1 core subunits (**Figure 1C - E**).

The AlphaFold3 model predicted that the Scm FCS finger domain (Scm_FCS_, Scm[Gly55-Ser135]) forms an extensive interface with the Pcl Tudor domain (Pcl_Tudor_ Pcl[Pro347-Leu399]), involving both hydrophobic and electrostatic interactions (**Figure 1F, Figure S1**). The interface is centred on a hydrophobic core formed by Scm Tyr75, Leu77, Pro78 and Ile92 and Pcl Phe358, Phe366, Leu368 and Leu399, flanked by electrostatic interactions between Scm Arg96 and Pcl Asp356 and Asp384 and between Scm Glu89 and Pcl Arg397 (**Figure 1F**).

### Reconstitution and biochemical validation of the Scm_FCS_–Pcl_Tudor_ interaction

To determine whether Scm_FCS_ and Pcl _Tudor_ interact directly, we expressed and purified the two domains individually (**Figure S2A, B**). Mixing purified Scm_FCS_ (Scm_55-135_) and Pcl _Tudor_ (Pcl_339-404_) resulted in formation of a stable complex that eluted as a single peak of higher apparent molecular weight than either individual domain during size-exclusion chromatography (**Figure 2A**). SDS-PAGE analysis of the eluted fractions showed that both proteins co-eluted across the peak at an approximately 1:1 ratio (**Figure 2A**). Online buffer exchange native mass spectrometry (OBE-native MS) (VanAernum et al., 2020) confirmed formation of a Scm_FCS_– Pcl_Tudor_ complex with a 1:1 stoichiometry (**Figure 2B**).

**FIGURE 2.**
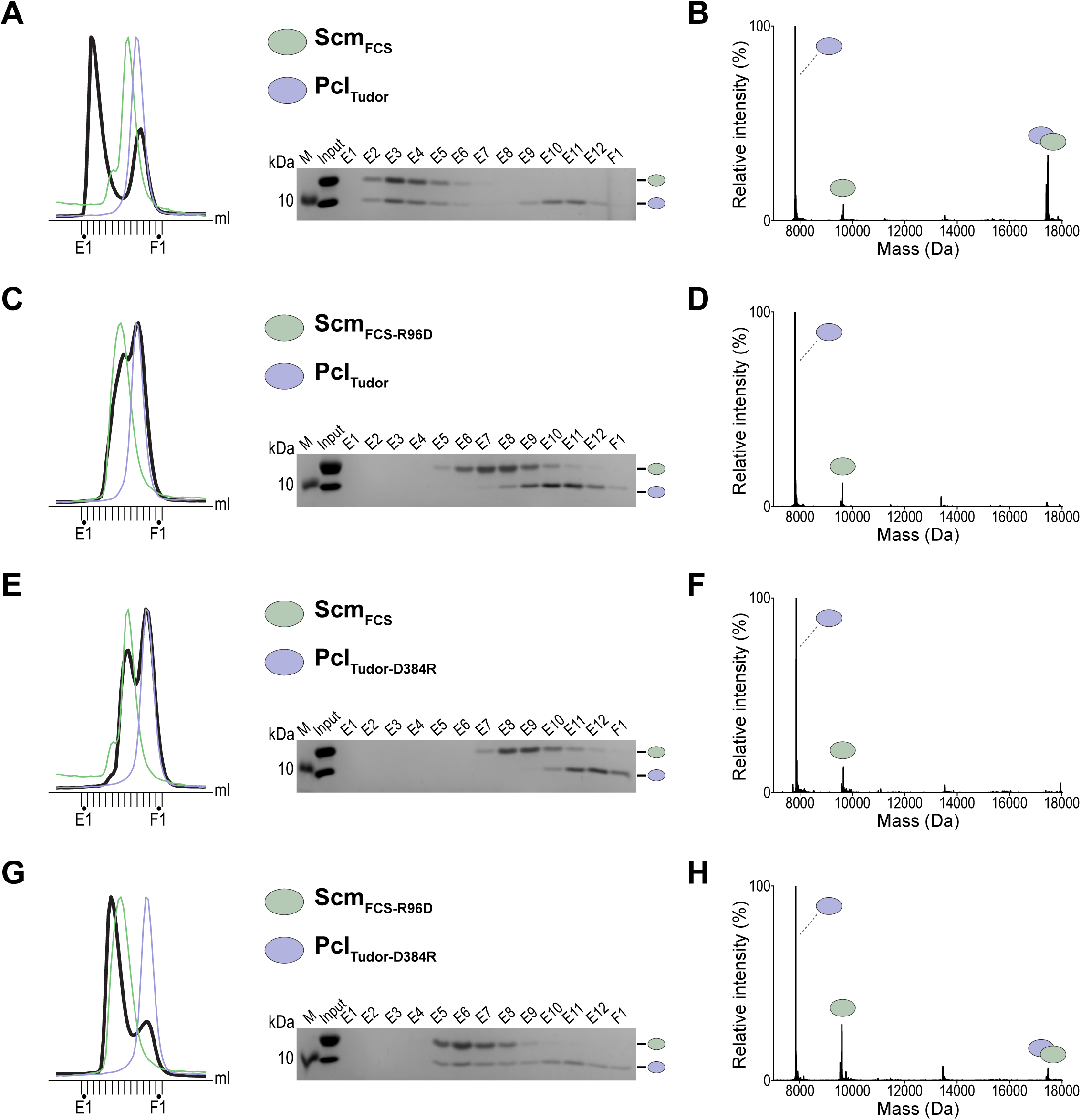
Biochemical validation of the Scm FCS finger–Pcl Tudor domain interaction. **A**, **B**: Analysis of complex formation between wild-type Scm_FCS_ and Pcl_Tudor_ domains. **A**: Left, size-exclusion chromatography profile of the Scm_FCS_–Pcl_Tudor_ mixture (black), with profiles of the individual Scm_FCS_ (green) and Pcl_Tudor_ (light purple) domains (**Figure S2A**) shown for reference. Right, SDS-PAGE analysis of peak fractions from the Scm_FCS_–Pcl_Tudor_ mixture. **B**: OBE-native MS analysis of the Scm_FCS_–Pcl_Tudor_ mixture. **C**, **D**: Analysis of complex formation between Scm_FCS-R96D_ and wild-type Pcl_Tudor_. **C**: Left, size-exclusion chromatography profile of the Scm_FCS-R96D_–Pcl_Tudor_ mixture (black), with profiles of the individual Scm_FCS-R96D_ (green) and Pcl_Tudor_ (light purple) domains (**Figure S2A, B**) shown for reference. Right, SDS-PAGE analysis of peak fractions from the Scm_FCS-R96D_–Pcl_Tudor_ mixture. **D**: OBE-native MS analysis of the Scm_FCS-R96D_–Pcl_Tudor_ mixture. **E**, **F**: Analysis of complex formation between wild-type Scm_FCS_ and Pcl_Tudor-D384R_. **E**: Left, size-exclusion chromatography profile of the Scm_FCS_–Pcl_Tudor-D384R_ mixture (black), with profiles of the individual Scm_FCS_ (green) and Pcl_Tudor-D384R_ (light purple) domains (**Figure S2A, B**) shown for reference. Right, SDS-PAGE analysis of peak fractions from the Scm_FCS_–Pcl_Tudor-D384R_ mixture. **F**: OBE-native MS analysis of the Scm_FCS_–Pcl_Tudor-D384R_ mixture. **G**, **H**: Analysis of complex formation between Scm_FCS-R96D_ and Pcl_Tudor-D384R_. **G**: Left, size-exclusion chromatography profile of the Scm_FCS-R96D_–Pcl_Tudor-D384R_ mixture (black), with profiles of the individual Scm_FCS-R96D_ (green) and Pcl_Tudor-D384R_ (light purple) domains (**Figure S2B**) shown for reference. Right, SDS-PAGE analysis of peak fractions from the Scm_FCS-R96D_– Pcl_Tudor-D384R_ mixture. **H**: OBE-native MS analysis of the Scm_FCS-R96D_–Pcl_Tudor-D384R_ mixture.

To test the AlphaFold3 model experimentally, we targeted the electrostatic interface by introducing complementary charge-reversal mutations into Scm and Pcl. Substitution of Scm Arg96 with aspartate (Scm_FCS-R96D_) abolished complex formation with wild-type Pcl_Tudor,_ as assayed by size-exclusion chromatography with SDS-PAGE analysis of the eluted fractions, and by OBE-native MS (**Figure 2C, D**). Likewise, substitution of Pcl Asp384 with arginine (Pcl_Tudor-D384R_) prevented complex formation with wild-type Scm_FCS_ in both assays (**Figure 2E, F**). Thus, charge reversal of either interaction partner disrupted formation of the Scm_FCS_– Pcl_Tudor_ complex, consistent with the AlphaFold3-predicted interface.

To determine whether the predicted electrostatic interaction could be reconstituted, we mixed Scm_FCS-R96D_ with Pcl_Tudor-D384R_. Remarkably, the complementary charge-reversal mutations restored complex formation, yielding a stable Scm_FCS-R96D_-Pcl_Tudor-D384R_ complex that co-eluted during size-exclusion chromatography and was confirmed by OBE-native MS to form a 1:1 complex (**Figure 2G, H**). Rescue of complex formation by complementary charge-reversal mutations provides strong experimental validation of the AlphaFold3 model and demonstrates that the predicted Scm Arg96–Pcl Asp384 electrostatic interaction is a key determinant of the Scm–Pcl interface. Together, these findings provide the molecular basis for the previously observed selective association between canonical PRC1 and PRC2.1 and identify direct Scm–Pcl binding as the physical link between the two complexes.

## DISCUSSION

This study provides two advances. First, we developed a native affinity-purification strategy based on cryo-milling and low-salt extraction that recovers higher-order Polycomb assemblies from *Drosophila* embryos. Second, we identified and biochemically validated a direct interaction between the Scm FCS finger and the Pcl Tudor domain as the molecular basis for the selective association between canonical PRC1 and PRC2.1 in *Drosophila*. These findings provide a molecular explanation for the selective association between canonical PRC1 and PRC2.1 previously observed in cross-linking-based affinity purification studies (Kang et al., 2022; Kang et al., 2015). Beyond establishing this molecular mechanism, our findings raise both technical and biological implications.

A technical aspect worth highlighting is that the native purification strategy established here not only recovered canonical PRC1 together with co-associated PhoRC and PRC2.1, which are co-bound with PRC1 at PREs, but also resulted in co-isolation of the O-GlcNAc transferase Sxc/Ogt. O-GlcNAc modification of Ph by Sxc/Ogt at a serine/threonine-rich region is required to prevent Ph aggregation (Gambetta & Müller, 2014; Gambetta et al., 2009). Although we did not investigate the interaction between Sxc/Ogt and Ph further, recovery of the modifying enzyme in Ph purifications suggests that the native purification strategy may preserve transient enzyme–substrate interactions that are normally disrupted during conventional biochemical purification. One speculative possibility is that the purification captured Sxc/Ogt while engaged in modification of Ph.

A biological implication of our findings is that the physical association between canonical PRC1 and PRC2.1 is mediated by the accessory subunits Scm and Pcl rather than by the conserved core subunits of the two complexes. Together with previous structural studies (Alfieri et al., 2013; Frey et al., 2016; Kim et al., 2005), our work establishes Scm as a central interaction hub that links canonical PRC1 and PRC2.1 within PRE-associated Polycomb complex assemblies. Scm is anchored to PRE-bound PhoRC through SAM–SAM interactions with the PhoRC subunit Sfmbt, while the EH surface of the Scm SAM domain engages the SAM domain of Polyhomeotic, thereby connecting canonical PRC1 to PRE-bound PhoRC (Frey et al., 2016; Kim et al., 2005). We identify here the N-terminal FCS finger of Scm as a second interaction interface that binds the Tudor domain of Pcl, thereby providing the physical link between canonical PRC1 and PRC2.1. Together, these interactions provide a molecular framework for the assembly and interconnection of distinct Polycomb protein complexes at PREs.

Notably, Scm and Pcl have both diverged substantially during evolution. Mammalian SCMH1 and SCML2 retain the conserved MBT and SAM domains but lack the N-terminal FCS finger (Bonasio et al., 2010) that we identify here as the Pcl-binding module in *Drosophila* Scm. Conversely, mammalian PCL proteins contain canonical Tudor domains that recognize H3K36me3 through an aromatic cage (Ballaré et al., 2012; Musselman et al., 2012), whereas the Tudor domain of *Drosophila* Pcl is atypical and lacks methyl-lysine binding activity (Friberg et al., 2010). The Scm–Pcl interaction described here therefore is unlikely to be evolutionarily conserved in mammals. Instead, physical coupling between canonical PRC1 and PRC2.1 appears to have diverged substantially during evolution and, if it exists in vertebrates, is likely to be mediated by a distinct molecular mechanism.

In an independent study published after completion of this manuscript, Lyons et al. (2026) also identified a direct interaction between the Scm FCS finger and the Pcl Tudor domain and investigated its role in PRC2.1 recruitment to Polycomb bodies and Polycomb repression *in vivo*.

## MATERIALS AND METHODS

### Affinity purification from cryo-milled embryos

2–16-hour old embryos expressing NTAP-tagged Ph-p_1-1589_ (Frey et al., 2016) and Oregon-R control embryos were dechorionated, washed thoroughly, snap-frozen in liquid nitrogen and stored at -80 °C. Frozen embryos were transferred to a cryogenic grinder (SPEX SamplePrep Freezer/Mill 6875) and ground in liquid nitrogen at 15 Hz (cycles per second) for six cycles with 2 min run time each, and 1-minute cooling time in between cycles. The embryo powder was stored at -80 °C until use.

For each condition, affinity purification was performed in triplicates as follow. 0.8 g of cryo-milled material was resuspended in 3.2 ml of lysis buffer P50 (50 mM potassium phosphate pH 8.0 and 0.1% NP-40), supplemented with 1x protease inhibitor cocktail (Roche, 04693132001) and 1 mM dithiothreitol (DTT). Once a homogenized lysate was obtained, it was equally divided into three and each fraction was centrifuged at 18,000 g for 15 min at 4 °C to remove cell debris and lipids. Cleared lysates were incubated with 25 μl of protein G Dynabeads (Invitrogen, 10004D) pre-coupled to anti-protein A IgG (Sigma-Aldrich P2921) for 30 min at 4 °C with gentle rotation. Beads were washed three times with P50 buffer prior to mass spectrometry analysis.

### LC-MS/MS and differential protein abundance analyses

#### Sample preparation

A total of 100 µL of SDC buffer containing 1% sodium deoxycholate (SDC), 40 mM 2-chloroacetamide (Sigma-Aldrich), and 10 mM tris(2-carboxyethyl)phosphine (TCEP; Thermo Fisher Scientific) in 100 mM Tris-HCl (pH 8.0) was added to the magnetic beads and incubated for 20 min at 37 °C. The samples were subsequently diluted 1:1 with water. Proteins were digested with 0.5 µg LysC for 1.5 h at 37 °C, followed by overnight digestion with 0.5 µg trypsin (Promega) at 37 °C. Peptides in the supernatant were separated from the beads using a magnetic rack. The peptide solution was acidified to a final concentration of 1% trifluoroacetic acid (TFA; Merck) and purified using SCX StageTips. Approximately 200 ng of peptides was subsequently loaded onto Evotips (Evotip Pure, Evosep).

#### LC-MS measurement

Peptides were eluted from the Evotips onto a 15 cm PepSep C18 column (15 cm × 150 µm, 1.5 µm; Bruker Daltonics) using an Evosep Eno HPLC system (Evosep). The column temperature was maintained at 50 °C, and peptide separation was performed using the 30 samples-per-day (SPD) method. Mass spectrometric data were acquired on a timsTOF Pro mass spectrometer controlled by timsControl software and operated in data-independent acquisition parallel accumulation–serial fragmentation (DIA-PASEF) mode. The MS scan range was set to 100– 1700 m/z, with an ion mobility range of 1/K₀ = 0.70–1.30 Vs cm⁻². Equal ion accumulation and ramp times of 100 ms were used in the dual TIMS analyzer, resulting in a spectral acquisition rate of 9.52 Hz. DIA-PASEF scans covered the 350.2–1199.9 Da range using 42 DIA-PASEF windows, each assigned to a single TIMS scan. Precursor isolation windows were alternated, resulting in a total cycle time of 2.21 s. Collision energy was linearly ramped from 45 eV at 1/K₀ = 1.30 Vs cm⁻² to 27 eV at 1/K₀ = 0.85 Vs cm⁻².

#### Data analysis

Raw data were processed using Spectronaut version 20.2 (Biognosys) in directDIA+ (library-free) mode. Spectra were searched against the UniProt database of Drosophila (SwissProt and TrEMBL, downloaded in 2023). Carbamidomethylation of cysteine residues was specified as a fixed modification, whereas methionine oxidation and protein N-terminal acetylation were included as variable modifications. Protein abundances across samples were quantified using label-free quantification (LFQ; MaxLFQ) at the MS2 level.

For differential protein abundance analysis, proDA (version 1.20.0) (Ahlmann-Eltze & Anders, 2020) method was performed by using log_2_ transformed iBAQ (intensity-based absolute quantification) values as input. Pairwise comparison was conducted between NTAP-tagged Ph-p purification and mock purification from Oregon-R embryonic extracts. Protein abundance in NTAP-Ph-p pull-down was calculated from the average abundance across samples (for the NTAP-Ph-p and the mock pull-down) and the fold change between conditions, as provided by proDA. Only nuclear proteins were included for the analysis according to a pre-defined classification based on UniProtKB taxonomy ID 7227 (*Drosophila melanogaster*) and Gene Ontology term GO: 0005634 (nucleus). Significance is defined by setting adjusted p-value lower than 0.1 and fold change between NTAP-Ph-p and mock purifications higher than 4.

#### Protein expression constructs

*Drosophila melanogaster* Scm_FCS_ (Scm[55-135]) and Pcl_Tudor_ (Pcl[339-404]) were cloned separately into modified monocistronic pEC-A-3C-GST *Escherichia coli* expression vectors using ligation-independent cloning system. This vector encodes an N-terminal 6xHis-GST tag followed by an HRV 3C cleavage site. UniProt accession numbers for Scm and Pcl are Q9VHA0 and Q24459, respectively. Scm_FCS-R96D_ and Pcl_Tudor-D384R_ mutant expression constructs were generated by standard site-directed mutagenesis.

#### Protein expression and purification

Recombinant Scm_FCS_, Scm_FCS-R96D_, Pcl_Tudor_ and Pcl_Tudor-D384R_ were separately expressed in *E. coli* Rosetta (DE3) pLysS cells. Transformed cells were grown at 37 °C in 10xP TB to an OD_600_ of approximately 1.0. Protein expression was subsequently induced by the addition of 0.5 mM isopropyl β-D-1-thiogalactopyranoside overnight at 18 °C. Cells were harvested and lysed by sonication, and the recombinant proteins were purified using Ni-NTA affinity chromatography.

Affinity tags were cleaved with HRV 3C protease (PreScission protease; MPI of Biochemistry Protein Core Facility), during dialysis against 25 mM Tris-HCl (pH 7.5), 250 mM NaCl, 10% glycerol overnight at 4 °C. The samples were subjected to reverse Ni-NTA affinity chromatography to remove affinity tags. The resulting flow through fractions were then further purified by size-exclusion chromatography (SEC) in a buffer containing 20 mM Tris-HCl (pH 7.5), 250 mM NaCl, 10% glycerol, and 2 mM DTT. The purity of each recombinant protein was assessed on polyacrylamide gels. Purified proteins were snap-frozen and stored at -80 °C until use.

#### Size-exclusion chromatography analysis

To assess the interaction between Scm_FCS_ and Pcl_Tudor_, analytical SEC was performed. Four mixtures were prepared containing either wild-type or mutant recombinant proteins as follows: Scm_FCS_ and Pcl_Tudor_, Scm_FCS-R96D_ and Pcl_Tudor_, Scm_FCS_ and Pcl_Tudor-D384R_, and Scm_FCS-R96D_ and Pcl_Tudor-D384R_. The protein mixtures were incubated for approximately 30 min at 4 °C. The samples were then injected into a Superdex 75 increase 3.2/300 column (ÄKTApurifier 10, Cytiva) at a flow rate of 0.03 ml/min and with a fractionation volume of 0.025ml. SEC was performed at 4 °C in a buffer containing 20 mM Tris-HCl (pH 7.5), 250 mM NaCl, and 2 mM DTT. Individual recombinant proteins were analyzed as controls under the same conditions with same injection loop and running buffer. The eluted fractions – E1 to F1 for mixtures, and peak fractions for individual proteins – were collected and analyzed by SDS-PAGE.

### OBE native mass spectrometry

#### Sample preparation and online buffer exchange

Four mixtures were prepared as described in size-exclusion chromatography analysis section. Online buffer exchange coupled to native MS was performed using a Thermo Scientific Vanquish Flex UHPLC system with duo pumps coupled to a Thermo Scientific Orbitrap Exploris 480 with BioPharma option. Online buffer exchange used a Thermo Scientific NativePac OBE-1 SEC column. The mobile phase was 200 mM ammonium acetate, run isocratically at 0.2 mL/min over a 10min method. Buffer-exchanged proteins eluted first and were directed to the MS; non-volatile salts eluted later and were diverted to waste using a six-port valve.

#### Native MS measurement

Native MS analyses were performed on a Thermo Scientific Orbitrap Exploris 480 with BioPharma option, operated in Intact Protein application mode. The spray voltage was set to 3.4 kV (positive ion, HESI source), the RF lens to 150% and sheath gas to 15. The ion transfer tube temperature was 280 °C. Source fragmentation was set to 60 V. Data were acquired in profile mode at a resolution setting of 15,000 at m/z 200 with transient averaging enabled. The scan range was m/z 1000–8000. The AGC target was set to Custom at 300%, with a maximum injection time of 200 ms and one microscan per spectrum.

#### Data analysis

Spectra were averaged over the retention-time window of the buffer-exchanged protein peak (2.10–2.35 min), selected to exclude a later-eluting polymer contaminant, and deconvolved using UniDec v8.2.1.

Deconvolution used a mass range of 7,000–18,000 Da sampled every 0.1 Da, a charge range of 3–20 (observed charge states 4–8), and an m/z range of 990–3600. The peak shape was modelled as a Gaussian with a fixed peak FWHM (mzsig) of 1.0 Th; charge smooth width and point smooth width were both 1.0 and mass smooth width was 0. The number of iterations was 50. A curved background subtraction (buffer 100) and an intensity threshold of 0.01 were applied during data processing. The adduct mass was set to the mass of a proton (1.007276 Da).

Peak detection used a peak window of 5.0 Da and a peak detection threshold of 0.02, with a peak plotting threshold of 0.1 and a minimum peak separation of 0.025. Harmonic suppression was enabled and the native charge offset was constrained to ±3 to suppress spurious assignments at integer multiples of the monomer mass.

Species were assigned using the UniDec Oligomer and Mass Tools with a matching tolerance of 5 Da, taking Zn²⁺ binding as +63.36 Da (Zn − 2H). Peaks supported by fewer than two charge states or with a charge-distribution standard deviation below 0.30, were classified as deconvolution artifacts and excluded from the reported species.

## ACKNOWLEDGMENTS

We thank Sven Schkölziger for help with protein expression, Maria Victoria Sanchez Caballero for help with mass spectrometry analysis (RRID:SCR_025745), Fabien Bonneau for help with generating AlphaFold3 predictions for inputs exceeding the standard 5,000-token limit, Anastasiia Gonchar for advice on cryo-milling of embryos and Christian Benda and Maria Ciapponi for discussions. This work was supported by the MPG.

## AUTHOR CONTRIBUTIONS

B.Y., J.B. and J.M. conceived the project. B.Y. designed, performed, and analyzed the experiments and prepared the figures. J.B. performed the initial AlphaFold3 analyses that identified the Scm–Pcl interaction. B.S. performed the mass spectrometry analyses. J.M. wrote the manuscript with input from all authors.

## CONFLICT OF INTEREST STATEMENT

The authors declare no competing or financial interests. The funder had no role in study design, data collection and analysis, decision to publish or preparation of the manuscript.

## DATA AVAILABILITY STATEMENT

The native mass spectrometry and proteomics data have been deposited at the ProteomeXchange Consortium via the PRIDE partner repository with the dataset identifier PXD084488.

## SUPPLEMENTARY MATERIAL

**Figure S1.**
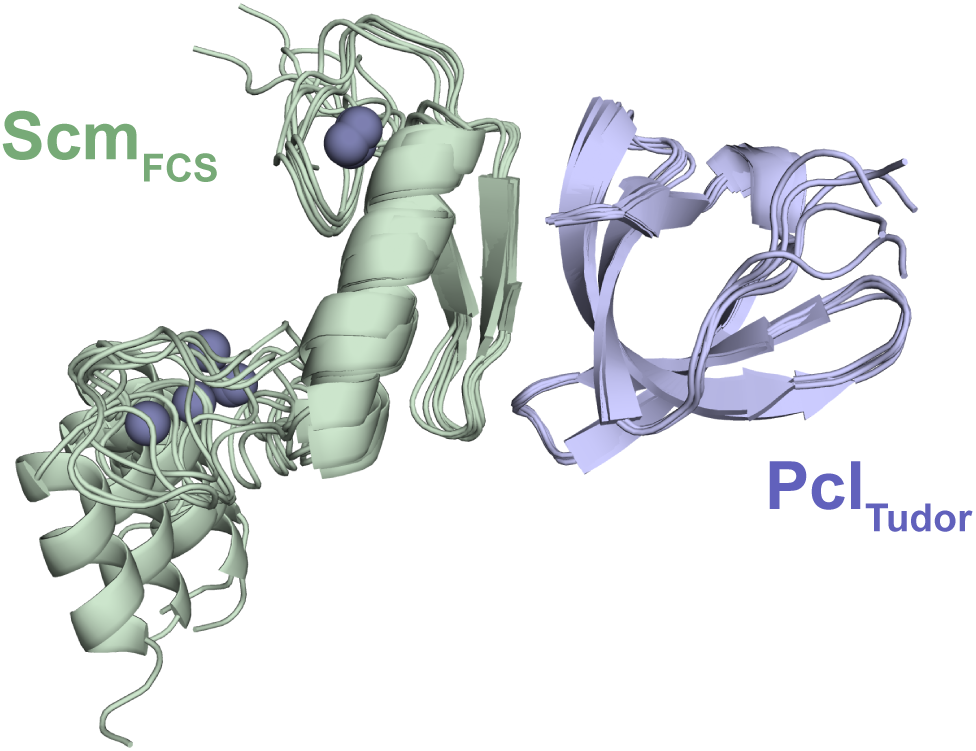
Superposition of AlphaFold3 models of the Scm FCS finger–Pcl Tudor domain complex. Superposition of the five AlphaFold3 models of the Scm FCS finger (ScmFCS)–Pcl Tudor domain (PclTudor) complex.

**Figure S2.**
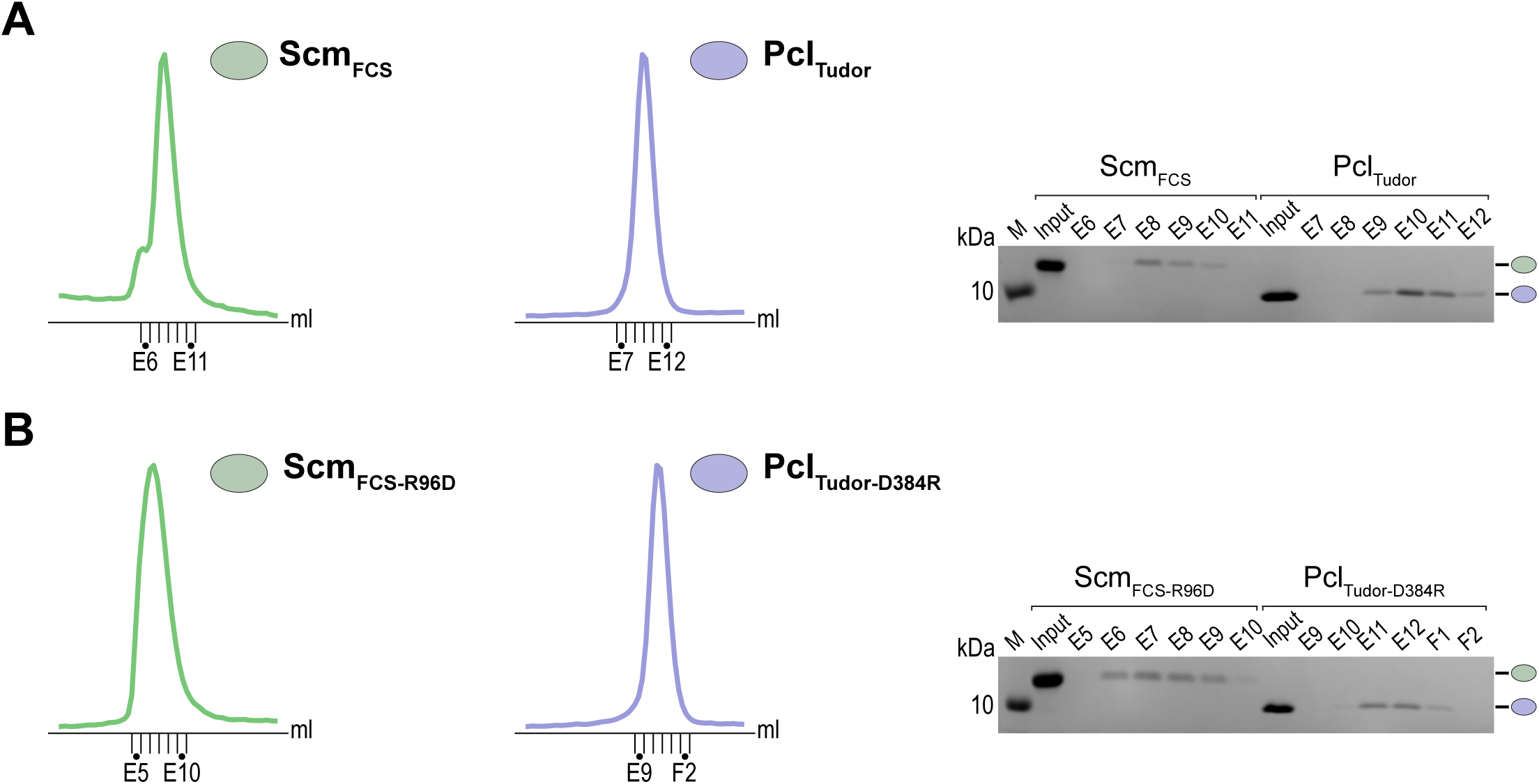
Purification of recombinant Scm FCS finger and Pcl Tudor domains. **A**: Size-exclusion chromatography and SDS-PAGE analysis of purified wild-type ScmFCS and PclTudor. **B**: Size-exclusion chromatography and SDS-PAGE analysis of purified Scm_FCS-R96D_ and Pcl_Tudor-D384R_.

## Notes

### Competing Interest Statement

The authors have declared no competing interest.

